# Multicenter validation of a machine learning algorithm for 48 hour all-cause mortality prediction

**DOI:** 10.1101/427054

**Authors:** Hamid Mohamadlou, Saarang Panchavati, Jacob Calvert, Anna Lynn-Palevsky, Christopher Barton, Grant Fletcher, Lisa Shieh, Philip B Stark, Uli Chettipally, David Shimabukuro, Mitchell Feldman, Ritankar Das

**Affiliations:** Dascena, Inc., Hayward, CA, United States; Department of Emergency Medicine, University of California San Francisco, San Francisco, CA, United States; Division of Internal Medicine, University of Washington School of Medicine, Seattle, WA, United States; Department of Medicine, Stanford University School of Medicine, Stanford, CA, United States; Department of Statistics, University of California Berkeley, Berkeley, CA, United States; Kaiser Permanente South San Francisco Medical Center, South San Francisco, CA, United States; Department of Anesthesia and Perioperative Care, University of California San Francisco, San Francisco, CA, United States; Division of General Internal Medicine, Department of Medicine, University of California San Francisco, San Francisco, CA, United States

**Keywords:** Machine learning, mortality, electronic health record, prediction

## Abstract

**Purpose:** This study evaluates a machine-learning-based mortality prediction tool.

**Materials and Methods:** We conducted a retrospective study with data drawn from three academic health centers. Inpatients of at least 18 years of age and with at least one observation of each vital sign were included. Predictions were made at 12, 24, and 48 hours before death. Models fit to training data from each institution were evaluated on hold-out test data from the same institution and data from the remaining institutions. Predictions were compared to those of qSOFA and MEWS using area under the receiver operating characteristic curve (AUROC).

**Results:** For training and testing on data from a single institution, machine learning predictions averaged AUROCs of 0.97, 0.96, and 0.95 across institutional test sets for 12-, 24-, and 48-hour predictions, respectively. When trained and tested on data from different hospitals, the algorithm achieved AUROC up to 0.95, 0.93, and 0.91, for 12-, 24-, and 48-hour predictions, respectively. MEWS and qSOFA had average 48-hour AUROCs of 0.86 and 0.82, respectively.

**Conclusion:** This algorithm may help identify patients in need of increased levels of clinical care.

## Introduction

Timely identification of patients with elevated risk for in-hospital mortality is necessary to best allocate limited and costly hospital resources, focus care to prevent patients from deteriorating, and anticipate probable patient outcomes [1]. Appropriate determination of which patients require ICU admission is especially important for appropriate resource allocation: although ICU beds account for less than 10% of beds in United States hospitals, ICU bed use and associated costs continue to rise, nearly doubling between 2000 and 2010 [2, 3]. Advance prediction of deterioration and death can alert medical teams to the need for more aggressive care while also helping to minimize overtreatment of more stable patients, in turn lowering health care costs. Mortality prediction tools that are accurate at long lookahead times have particular promise for providing optimal care and allocating hospital resources [4, 5].

There are several existing mortality prediction tools; these systems generally use rules-based approaches. Such rules-based systems include the Acute Physiology, Age, Chronic Health Evaluation (*APACHE*) [6], the Modified Early Warning Score (*MEWS*) [7], the Sepsis-Related Organ Failure Assessment (*SOFA*) [8], and the quick SOFA (*qSOFA*) score [9], but the utility of such tools in clinical settings is limited due to inadequate specificity and sensitivity [10, 11]. These scores use a collection of baseline patient characteristics, assign a linear weight to each characteristic, and sum the weighted scores to create an overall risk factor. Because these tools are based on a single snapshot of a patient’s characteristics, they cannot incorporate trends in patient conditions, which can be useful for predicting patient decline [12]. Changes in scores may be informative, but a tool that directly incorporates trends in patient vital signs may be more reliable.

A machine learning mortality prediction tool that is integrated into an electronic health record (EHR) system can take such trends into account. Machine learning algorithms can produce a score that depends not only on linear combinations of the input variables, but also on trends in those variables. Previous studies have demonstrated that machine learning mortality risk scores that incorporate temporal information can more accurately predict patient outcomes [12–14]. Moreover, machine learning algorithms are readily optimized for different populations, while existing rule-based methods are “one size fits all” [13].

We have developed a machine learning mortality risk prediction tool with a 48-hour prediction horizon using only six vital signs and Glasgow Coma Scale (GCS) values as input. In this investigation, we have demonstrated improved performance over the existing methods, MEWS and qSOFA. We validated predictions across datasets from three academic health centers in the United States, to examine the robustness of the predictions across patient populations in different hospitals without customized training.

## Materials and Methods

### Data Sources

Patient records were collected by Stanford Medical Center in Stanford, California; the University of California, San Francisco Medical Center in San Francisco, CA (UCSF); and University of Washington Medical Center (UW) in Seattle, Washington. Data from Stanford included records from 515,452 inpatients from all hospital wards from December 2008 to May 2017. Data collected at UCSF contained information on 95,869 patients (both inpatients and outpatients) across all hospital wards from the Mount Zion, Mission Bay, and Parnassus Heights medical campuses. We used inpatient data from June 2011 to March 2016 drawn from patient EHR charts. Data from UW included records from 32,936 adult patients from all hospital wards from January 2014 to March 2017.

Data collection was passive and had no impact on patient safety. All data were deidentified in compliance with the Health Insurance Portability and Accountability Act (HIPAA) Privacy Rule. Studies performed on the de-identified data constitute non-human subject studies, and therefore our study did not require Institutional Review Board approval.

### Data Processing

For all three data sources, we included only records for patients aged 18 or older, who had at least one recorded observation of each required measurement (heart rate, respiratory rate, peripheral oxygen saturation (SpO_2_), temperature, systolic blood pressure, diastolic blood pressure, and GCS). Additionally, we filtered-out patient records for which there were no raw data or no discharge or death dates. This resulted in 31,292 patients from Stanford, 47,289 patients from UCSF, and 32,878 patients from UW.

Hourly measurements were required for each input variable; if no measurement was available for a given hour, a carry-forward method was used to impute values from the most recent past measurement. For some patients with infrequent measurements of one or more vital signs (not including GCS), this simple imputation resulted in many consecutive hours with identical values, which we quantified for each vital sign as the fraction of measurements which were equal to the mode of the measurements. As distinct vital sign measurements are unlikely to be identical and because the mode only counts identical values, if this mode fraction is high, it indicates that most of the measurements of the vital sign have been imputed. We excluded patients with two or more vital signs which had mode fractions exceeding 0.90. After discarding patients with multiple, high mode fractions, there were 24,614 patients from Stanford, 46,980 patients remaining from UCSF, and 32,718 patients from UW. Supplementary Table 1 lists the number of patients sequentially meeting each inclusion criterion.

### Gold Standard

The outcome of interest was in-hospital patient mortality, which was determined retrospectively for each patient. In the Stanford and UW datasets, in-hospital mortality was indicated by a death date field for each patient. In the UCSF dataset, we used the in-hospital mortality field for each patient. Consequently, positive-class labels were assigned to 1,810 patients of 46,980 from UCSF (3.85%), 263 patients of 24,614 from Stanford (1.07%), and 965 patients of 32,718 from UW (2.95%).

### The Machine Learning Algorithm

The classifier was created using the XGBoost method for creating gradient boosted trees. We applied the XGBoost package for Python [15] to the patient measurements and their temporal changes, where temporal changes included hourly differences between each measurement beginning three hours before prediction time. We minimally processed raw EHR data to generate features. Following EHR data extraction and imputation, we obtained six hourly values for each of the seven measurement channels from that hour, and each hour prior up to three hours. We also calculated two difference values between the current hour and the prior hour, and between the prior hour and the hour before that. We concatenated these values from each measurement into a causal feature vector. Gradient boosting is an ensemble learning technique that combines results from multiple decision trees to create prediction scores. Tree branching is determined by the gradient boosting software, which discretized each feature into two possible categories, such as temperature above or below 100°F. The result of this branching brings a patient to the next node, where they may be categorized in a similar binary fashion based on the same or another clinical variable. A risk score is assigned to each terminal node of the resulting tree. We restricted tree depth to a maximum of six branch levels, set the learning rate parameter of XGBoost to 0.1, and restricted the tree ensembles to 200 trees to limit the computational burden.

To validate the boosted tree predictor when training and testing was performed on data from the same institution, we used three-fold cross validation. For each model, two thirds of the data set was randomly selected as a training set and one third was retained as an independent test set. Performance was evaluated on the independent testing set. Reported performance metrics are the average performance of 20 separately trained models, where each model was trained on shuffled portions of the data and tested on the remaining disjoint third. We ran the boosted tree ensemble for mortality prediction at 12, 24, and 48 hours before death in order to evaluate its performance as time before patient death increased. For negative class examples, because there was no mortality event from which a specific, applicable time-point could be computed for survivors, we used their time of discharge instead. Models were independently trained for each distinct lookahead time.

### Comparison to Rules-Based Methods

To calculate AUROC values for the rules-based predictors, we calculated MEWS and qSOFA score for patients in the UCSF database. MEWS and qSOFA scores were calculated on the entire dataset. We calculated the MEWS score using systolic blood pressure, heart rate, respiratory rate, temperature, and GCS values extracted from EHR data. Scores were calculated using the system described in Fullerton et al [16]. MEWS is often calculated from Alert, Voice, Pain, Unresponsive (AVPU) values; however, the datasets used in this experiment more reliably contained GCS data. We therefore converted GCS to AVPU values as follows: 13–15 GCS points as Alert; 9–12 GCS points as Voice; 4–8 GCS points as Pain; ≤ 3 GCS points as Unresponsive. This conversion is similar to the one described in Kelly et al. [17], but eliminates the overlap between GCS ranges. Such conversions have been utilized in past retrospective studies [18] under similar data availability conditions and do not appear to lower the predictive accuracy of MEWS. The qSOFA score values were calculated with blood pressure, respiratory rate, and GCS values. To compare qSOFA and MEWS to the boosted tree predictor, the boosted tree predictor was trained and tested on UCSF data using all six clinical vital signs and GCS.

### Cross Population Validation Experiments

To test the performance of the boosted tree predictor when subjects in the training data differ demographically and clinically from those in the test data, we performed cross-population experiments. We trained the boosted ensemble for mortality prediction exclusively on one of the three datasets, using the entire dataset for training, and then tested model performance on the remaining two datasets without any site-specific retraining. Testing was performed on the entire dataset for each non-training hospital. We performed these experiments using only five patient vital signs: systolic blood pressure, diastolic blood pressure, heart rate, temperature, and respiratory rate. Other measurements were not included due to limited data availability in the Stanford Medical Center and University of Washington Medical Center datasets. For all cross-population experiments, the predictor was trained using XGBoost [15], in the same manner and using the same inputs as described above. We ran the cross population validation tests at 12, 24, and 48 hours before patient death.

## Results

The three data sources are all large academic hospitals, but differ in their patient demographics (Table 1). UCSF patients were typically older than patients at Stanford and UW. 25.92% of UCSF patients were over 70 years old. Approximately 43% of UCSF patients were between 50 and 70 years old, compared with approximately 34% of Stanford patients and 38% of UW patients. Differences in patient demographics allowed us to test whether the algorithm’s predictions are robust across different patient populations.

**Table 1:**
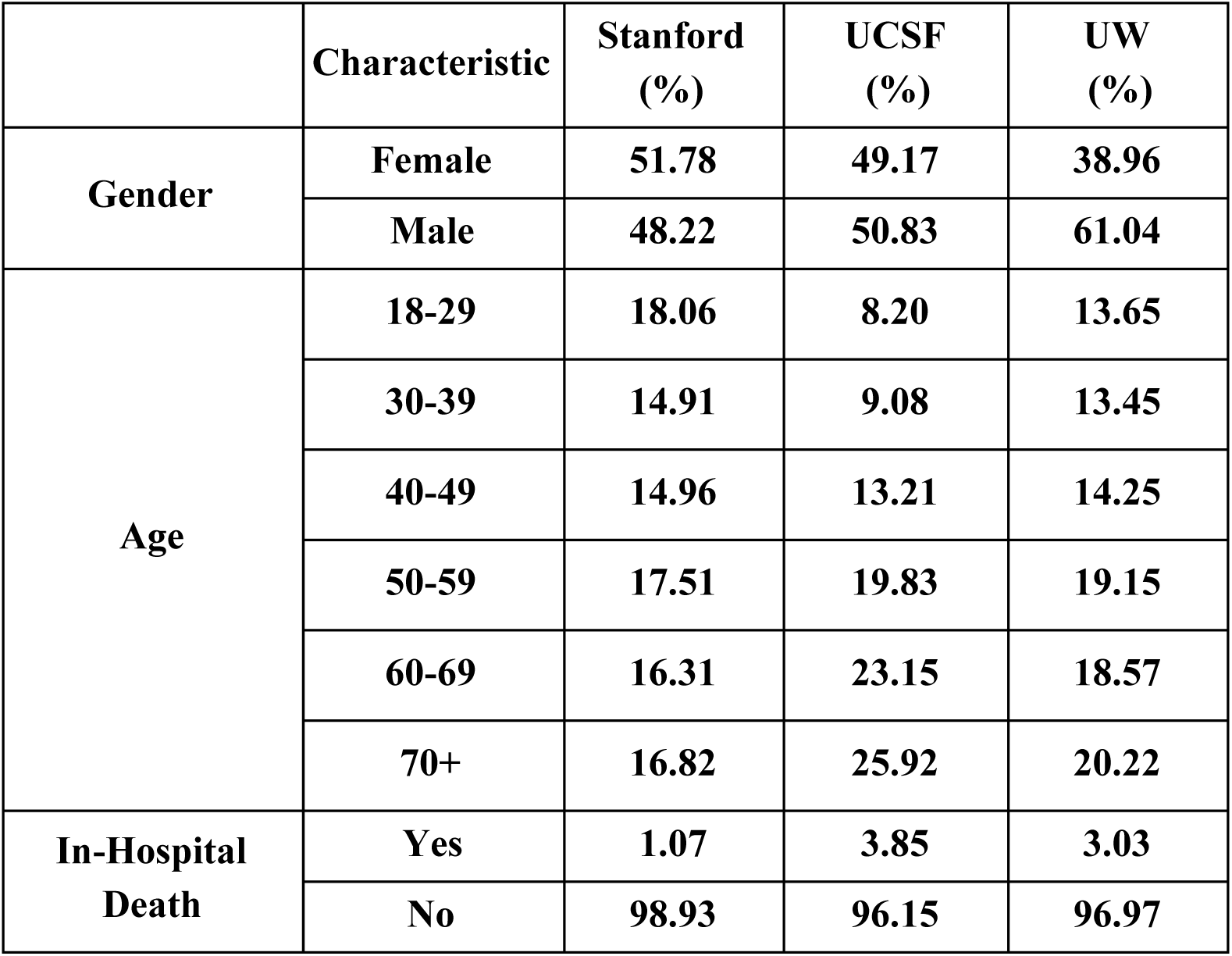
Patient demographic information for processed Stanford, University of California, San Francisco (UCSF), and University of Washington (UW) cohorts.

|  | **Characteristic** | **Stanford (%)** | **UCSF (%)** | **UW (%)** |
| --- | --- | --- | --- | --- |
| **Gender** | **Female** | **51.78** | **49.17** | **38.96** |
|  | **Male** | **48.22** | **50.83** | **61.04** |
| **Age** | **18-29** | **18.06** | **8.20** | **13.65** |
|  | **30-39** | **14.91** | **9.08** | **13.45** |
|  | **40-49** | **14.96** | **13.21** | **14.25** |
|  | **50-59** | **17.51** | **19.83** | **19.15** |
|  | **60-69** | **16.31** | **23.15** | **18.57** |
|  | **70+** | **16.82** | **25.92** | **20.22** |
| **In-Hospital Death** | **Yes** | **1.07** | **3.85** | **3.03** |
|  | **No** | **98.93** | **96.15** | **96.97** |

The machine learning algorithm predicted patient mortality with a higher accuracy than qSOFA and MEWS for nearly all prediction windows on all datasets. When trained and tested on Stanford data, the MLA had AUROC of 0.95 for 48-hour mortality prediction (Figure 1A). When trained and tested on data collected from UCSF, the MLA had AUROC of 0.97 for 48 hour mortality prediction (Figure 1B). For training and testing on UW data, the MLA had AUROC of 0.93 for 48-hour prediction (Figure 1C).

**Figure 1:**
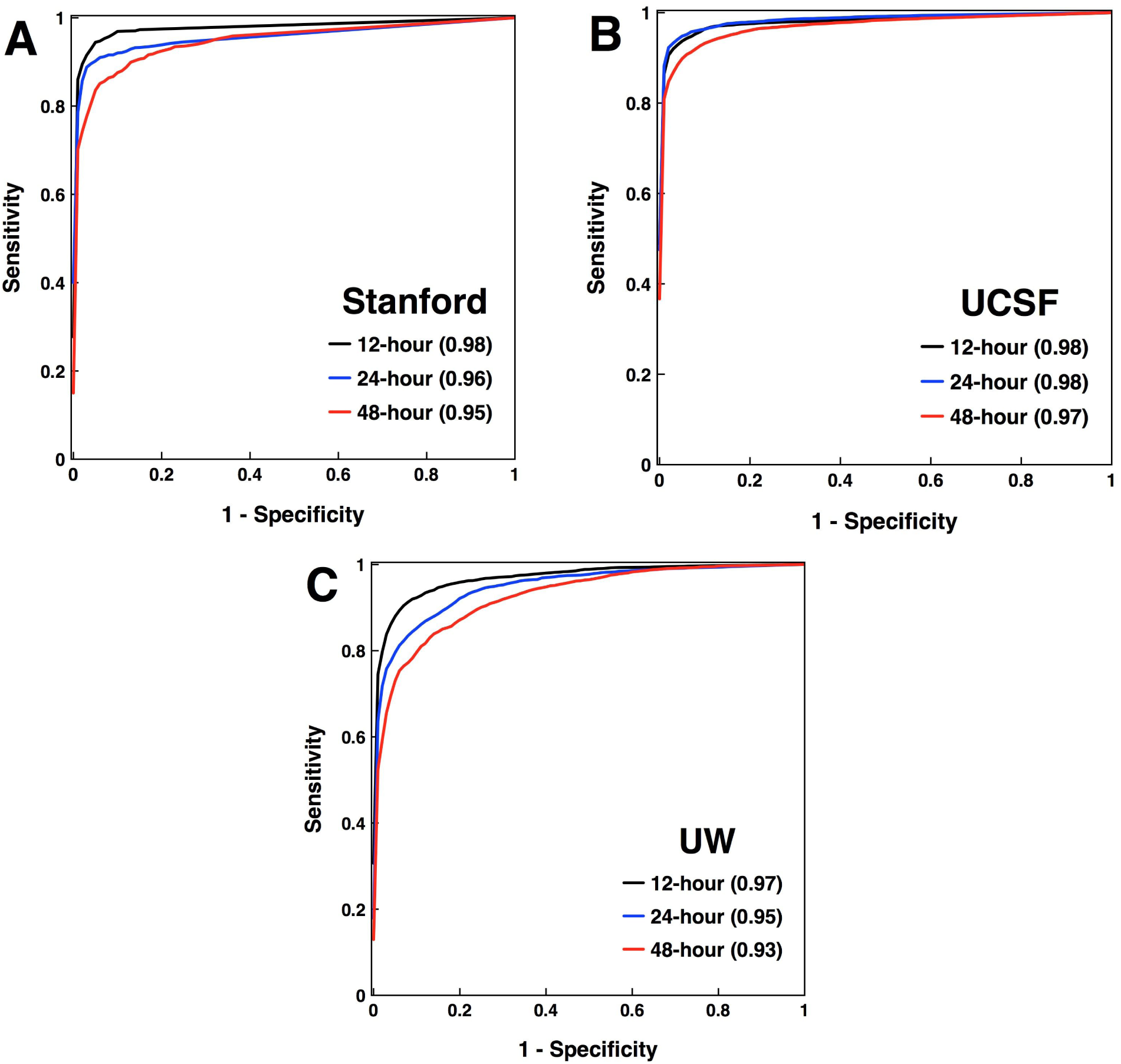
Comparison of the receiver operating characteristic (ROC) curve. ROC curves and area under the ROC (AUROC) for the machine learning algorithm are presented for 12-, 24-, and 48-hour mortality prediction with training and testing performed on A) **Stanford** patient data, B) **University of California, San Francisco (UCSF)** patient data, C) **University of Washington (UW)** patient data. The Stanford and UCSF predictions used systolic blood pressure, diastolic blood pressure, heart rate, temperature, respiratory rate, SpO_2_, and Glasgow Coma Scale, whereas the UW predictions used only systolic blood pressure, diastolic blood pressure, heart rate, temperature, and respiratory rate.

On the Stanford dataset, MEWS had an AUROC of 0.90 for 48-hour prediction, while the qSOFA score had an AUROC of 0.87 for 48-hour prediction (Table 2). On the UCSF dataset and for 48-hour prediction, MEWS and qSOFA had AUROCs of 0.93 and 0.89, respectively. On the UW dataset and for 48-hour prediction, MEWS and qSOFA had AUROCs of 0.75 and 0.70, respectively. Across the three datasets, MEWS and qSOFA had average AUROCs of 0.86 and 0.82, respectively. In comparison, the MLA had an average AUROC of 0.95 for 48-hour prediction.

For all prediction windows, the MLA had a significantly higher Diagnostic Odds Ratio (DOR) than either MEWS or qSOFA. Additional statistics (corresponding to specificity above 0.80, in the clinical operating range for the predictor) for the MLA trained on Stanford data, MEWS, and qSOFA, are presented in Table 2.

**Table 2:** Comparison of AUROC, Diagnostic Odds Ratio (DOR), sensitivity, and specificity values for the algorithm and qSOFA and MEWS scoring systems for mortality prediction. Predictions were performed **12, 24, and 48 hours in advance** patient death. All predictor training and testing was done on the Stanford data set using patient measurements for **heart rate, respiratory rate, temperature, SpO_2_, diastolic blood pressure, and systolic blood pressure and Glasgow Coma Scale (GCS)**. Each reported MLA value is the average of that value over 10-fold cross-validation. Consequently, the metrics have not been calculated directly from one another (e.g. MLA DOR does not agree with the ratio of LR+ to LR−).

|  |  | **Machine Learning Algorithm** | **MEWS Score** | **qSOFA Score** |
| --- | --- | --- | --- | --- |
| **12 hours before death** | **AUROC (95% CI)** | **0.979 (0.961, 0.997)** | 0.950 | 0.925 |
|  | DOR | 1945.21 | 101.73 | 85.73 |
|  | Sensitivity | 0.788 | 0.811 | 0.960 |
|  | Specificity | 0.994 | 0.960 | 0.782 |
|  | LR+ | 414.33 | 20.01 | 4.40 |
|  | LR- | 0.213 | 0.197 | 0.051 |
| **24 hours before death** | **AUROC (95% CI)** | **0.957 (0.945, 0.969)** | 0.928 | 0.891 |
|  | DOR | 598.64 | 57.02 | 35.22 |
|  | Sensitivity | 0.802 | 0.809 | 0.920 |
|  | Specificity | 0.989 | 0.966 | 0.753 |
|  | LR+ | 116.62 | 7.36 | 3.73 |
|  | LR- | 0.200 | 0.129 | 0.106 |
| **48 hours**<br>**before death** | **AUROC**<br>**(95% CI)** | **0.948**<br>**(0.925, 0.971)** | 0.895 | 0.865 |
|  | DOR | 180.23 | 29.96 | 22.72 |
|  | Sensitivity | 0.798 | 0.838 | 0.905 |
|  | Specificity | 0.960 | 0.853 | 0.705 |
|  | LR+ | 36.93 | 5.69 | 3.55 |
|  | LR- | 0.211 | 0.190 | 0.119 |

In the cross population experiments, the algorithm trained and tested on different data sets achieved AUROC values up to 0.95 at 12 hours before death, 0.93 at 24 hours before death and up to 0.91 at 48 hours preceding death (Table 3). Averaged across pairs of training and testing sets from different data sets, the AUROCs were 0.90, 0.86, and 0.84 for 12-, 24-, and 48-hour prediction. Across training sets, average 12-hour cross-population performance was best when training was done on the UW data set.

**Table 3:** Average AUROC values with 95% confidence intervals for each data set for cross-population training experiments. Confidence intervals were calculated from 10-fold cross-validation. Testing was performed **12, 24, and 48 hours in advance** of patient death. Risk scores were computed using **systolic blood pressure, diastolic blood pressure, heart rate, temperature, and respiratory rate**.

|  | UCSF<br>12 hrs<br>(95%<br>CI) | UCSF<br>24 hrs<br>(95%<br>CI) | UCSF<br>48 hrs<br>(95%<br>CI) | Stanford<br>12 hrs<br>(95%<br>CI) | Stanford<br>24 hrs<br>(95%<br>CI) | Stanford<br>48 hrs<br>(95%<br>CI) | UW<br>12 hrs<br>(95%<br>CI) | UW<br>24 hrs<br>(95%<br>CI) | UW<br>48 hrs<br>(95%<br>CI) |
| --- | --- | --- | --- | --- | --- | --- | --- | --- | --- |
| Trained<br>on<br>UCSF | **0.978**<br>(0.972,<br>0.983) | **0.979**<br>(0.977,<br>0.982) | **0.969**<br>(0.962,<br>0.975) | **0.919** | **0.870** | **0.833** | **0.916** | **0.908** | **0.862** |
| Trained<br>on<br>Stanford | **0.909** | **0.875** | **0.846** | **0.968**<br>(0.956,<br>0.981) | **0.919**<br>(0.890,<br>0.949) | **0.887**<br>(0.835,<br>0.938) | **0.815** | **0.810** | **0.768** |
| Trained<br>on<br>UW | **0.950** | **0.932** | **0.914** | **0.901** | **0.785** | **0.798** | **0.970**<br>(0.964,<br>0.976) | **0.948**<br>(0.935,<br>0.961) | **0.927**<br>(0.915,<br>0.939) |

## Discussion

When trained and tested on retrospective data, the boosted tree predictor more accurately predicted patient mortality than did the MEWS and qSOFA risk scoring systems, as evidenced by a battery of metrics at a particular operating point (Table 2) and by ROC curves, summarizing performance across operating points (Figure 1). We chose to present a comparison of our algorithm with the MEWS and qSOFA scoring systems because they are well-validated, easily measured prediction scores which are commonly used in for all-cause mortality across clinical settings in the United States.

In the present context, the AUROC can be somewhat misleading, as it considers performance at operating points which are not clinically relevant (e.g. sensitivity of 0.99 and specificity of 0.05) and because the low prevalence of in-hospital mortality in the three datasets allows even trivial predictors to obtain respectable accuracy (e.g. a predictor which guesses the patient will live, every time). For this reason, the comparison between the MLA, MEWS, and qSOFA in Table 2 is perhaps most informative. For 12-hour prediction, the differences in DOR and specificity are staggering; a positive-class prediction from the MLA at this operating point would substantially enrich one’s prior belief in a patient’s risk of in-hospital mortality.

Even when trained and tested on separate patient datasets, the boosted tree predictor maintained high levels of accuracy as demonstrated by AUROC values, up to two days in advance of patient death (Table 3). This comparison quantifies the robustness of the MLA to changes in patient populations, which are measurably different in terms of demographic information (Table 1) and which have different prevalences of in-hospital mortality. The UCSF and UW data sets were the most similar in these regards, as both sets tended to have older patients than the Stanford set, and had higher prevalences (3.85% and 2.95%, respectively) than in the Stanford set (1.07%). This is reflected in the cross-population results (Table 3), for which 12-hour prediction on the UW data set is much better when trained on the UCSF set than when trained on the Stanford set (respective AUROCs of 0.916 and 0.815).

Other machine learning-based methods have been developed for in-hospital mortality prediction for specific acute conditions such as sepsis [19] and cardiac arrest [20], or for specific settings such as the ICU [13]; however relatively little work has been done using machine learning methods to predict all-cause mortality across all hospital wards. A random forest model developed by Churpek et al. achieved an AUROC of 0.80 on hospital floor patients [21]. An ensemble learning approach by Pirracchio et al. reported an AUROC of 0.88 for in-hospital mortality in the ICU [22]. Taylor et al. describe a random forest model achieving up to 0.86 AUROC in the Emergency Department for patients with sepsis [23]. Escobar et al. have reported an in-hospital mortality AUROC up to 0.883 across all hospital wards using logistic regression [24]. However, each of these approaches requires extensive patient data including laboratory results, patient histories, and patient demographics. In contrast, the boosted tree predictor described here can make accurate mortality predictions using as few as five routinely collected vital signs.

While MEWS and qSOFA must be calculated with the same variables each time, the boosted tree predictor can use a variety of available inputs without sacrificing accurate patient mortality prediction. The boosted tree predictor can also be tailored to specific patient populations. This flexibility provides advantages over MEWS and qSOFA-based all-cause mortality predictions.

The boosted tree predictor’s combination of high sensitivity and specificity means that it can identify more at-risk patients than MEWS and qSOFA while also reducing the number of false alarms. The low specificity of many rules-based systems can lead to alarm fatigue; this desensitization to alarm systems can prevent early detection of diseases, and is recognized as a leading patient safety concern in the US [25]. The prediction system described here reduces this risk without sacrificing high sensitivity. The greater area under the curve within the clinical operating range of the boosted tree predictor as compared to MEWS and qSOFA further demonstrates the increased specificity of this predictor as compared to either rules-based score.

By providing more accurate warnings of mortality risk up to 48 hours in advance, this boosted tree predictor could provide clinical teams more time to intervene and potentially improve patient outcomes. Many conditions can be readily treated in their early stages but have high costs and mortality associated with their advanced stages. Survival rates for patients with septic shock patients have been shown to decrease by 7.6% each hour before antibiotics are administered after onset, and delays in acute kidney injury treatment can lead to total renal failure requiring kidney replacement, or to patient mortality [26, 27]. By predicting patient mortality up to two days in advance, outcomes of such patient could likely be improved.

The cross-population results also have promising clinical implications. Machine learning-based prediction systems must generally be trained on large amounts of retrospective data from a given hospital site, a process that is burdensome for the hospitals and can delay the implementation of life-saving systems. Because this prediction system maintained high accuracy when trained and tested on separate datasets, this predictor might outperform most mortality predictors even without site-specific training. This could allow hospitals to adopt the mortality predictor more rapidly and still improve patient outcomes.

There are several limitations to our study. Because this study was performed on retrospective data, we can not draw strong conclusions about its performance in a prospective clinical setting. Although there are important demographic differences across the datasets used in this study, all data came from large, urban research universities. Performance on a patient population that differs substantially from these, such as that of a rural hospital, may vary from this performance. Because of the retrospective nature of this work, we do not know how clinicians might adjust their actions based on risk scores. We also can not determine from this study whether more aggressive treatment would have prevented or postponed the deaths of those patients who died.

Reported performance metrics are the average performance of 10 separately trained models. This process was necessary due to the low incidence of mortality in our datasets; dividing the data into a larger number of folds for cross validation would provide too few examples of patient mortality for accurate training.

### Conclusion

The boosted tree predictor accurately predicted patient mortality 48 hours in advance of death using only patient vital signs. The algorithm demonstrated significantly improved accuracy over two commonly used mortality risk stratification tools. In future studies, we intend to test the algorithm prospectively using real-time clinical data. In a clinical setting, this algorithm may help clinicians identify those patients for whom more intensive care would prevent deterioration.

## Acknowledgements

We gratefully acknowledge Jana Hoffman and Emily Huynh for their suggestions and assistance in editing this manuscript. We also thank Thomas Desautels for his feedback during this study.

## Author Contributions

HM and RD conceived and designed this study. All authors contributed to acquisition, analysis, or interpretation of data. HM and ALP drafted the manuscript, and all authors revised the manuscript for important intellectual content. HM and PS performed statistical analysis. RD obtained funding.

